# Evolutionary analysis of vertebrate KCNH voltage-gated potassium channels and their expression in zebrafish embryos

**DOI:** 10.64898/2026.05.21.726828

**Authors:** Kuangyi Wu, Dingxun Wang, Ziyu Dong, Alice Yahui Zhou, GuangJun Zhang

**Author notes:** Corresponding author: GuangJun Zhang. These authors contributed equally to this work (Kuangyi Wu, Dingxun Wang).

## Abstract

Voltage-gated potassium channels (Kv) are a large family of potassium channels composed of 40 members across 12 subtypes. The *KCNH* genes encode 3 subfamilies of voltage-gated potassium channels: Kv10 (EAG, *ether à go go*), Kv11 (ERG, EAG-related gene), and Kv12 (ELK, EAG-like K). Kv channels play prominent roles in the neuronal and cardiovascular systems. Mutations in Kv channels have been linked to many human diseases, such as epilepsy, heart arrhythmias, and cancers. Significant progress has been made in understanding protein structures, physiological functions, and the pharmacological modifiers. However, the evolutionary history and gene expression of vertebrate *KCNH* genes during embryonic development remain largely unknown. We systematically identified and cloned 14 *kcnh* genes in zebrafish. Then, we examined vertebrate KCNH channel evolution by phylogenetic and syntenic analyses. Our data revealed that the three subtypes of *the KCNH* gene family have already evolved in invertebrates, long before the emergence of vertebrates. The number of vertebrate *KCNH* genes increased, most likely due to whole-genome duplications (WGDs). In addition, we examined zebrafish *kcnh* gene expression during early embryogenesis by *in situ* hybridization. Each subgroup’s genes showed similar but distinct gene expression domains with some exceptions. Most of them were expressed in neural tissues. Notably, *kcnh6a* showed robust expression in the developing heart, consistent with its conserved role in cardiac repolarization. Additionally, a few *kcnh* genes were transiently expressed in nonneural tissues, such as somites and the notochord, suggesting they may have a unique role in embryonic development. Our phylogenetic and developmental analyses of KCNH channels shed light on their evolutionary history and potential roles during embryogenesis, in line with their physiological functions and human channelopathies.

## INTRODUCTION

Potassium channels are aqueous viaducts within the cell membrane, though a few exceptions, such as TMEM175, can be found in organelles ^1^. All potassium channels selectively conduct potassium ions down their electrochemical gradient. Membrane potassium channels are evolutionarily conserved, and all possess a unique potassium-selective motif, (T/S)XGXGX, in the pore-forming loops (P-loop) ^2^. Based on protein structure and gating mechanisms, they are usually classified into four types: inwardly rectifying potassium channels (Kir), two-pore-domain potassium channels (K2P), Calcium-activated potassium channels (Kca), and voltage-gated potassium channels (Kv). The Kv channels are tetramers, and each α subunit contains six transmembrane (TM) domains: TM4 as a voltage sensor and TM5-6 as the walls of the potassium pore. The Kv channel family is the most diverse, comprising 40 channel genes that can be further classified into 12 subfamilies in the human genome ^3^.

The *KCNH* genes encode 3 subfamilies of voltage-gated potassium channels: Kv10 or EAG (ether à go go), Kv11 or ERG (EAG-related gene), and Kv12 or ELK (EAG-like K) channels ^4,5^. KCNH channels play essential roles in physiology and are critical for neuronal excitability ^6^, cardiac repolarization ^7^, and tumor cell growth ^8^. Many KCNH genes have been associated with human diseases. For example, KCNH1/Kv10.1 mutations were linked with Zimmermann–Laband syndrome and Temple–Baraitser syndrome ^9,10^. Both are rare disorders characterized by intellectual disability, facial dysmorphism, and abnormality of phalanges and nails. Mutations in human KCNH2/Kv11.1 cause epilepsy, Long QT-2 Syndrome, and short QT syndrome ^11,12^. Mutations of KCNH3/Kv12.2 and KCNH5/Kv10.2 were recently found to be associated with autism and epilepsy ^13,14^.

The KCNH channels have been extensively investigated, primarily for their roles in neurons and the heart. Their functions in other somatic tissues and in embryonic tissues are starting to be recognized. Recently, the well-known naturally occurring zebrafish longfin mutant, *lof*, was demystified ^15,16^. About 1Mb DNA fragment between *kcnh2a* and *prrx1a* genes on chromosome 2 is inverted and leads to the *kcnh2a* ectopic fin expression, which contributes to the elongated fins. Similarly, *kcnh8* was found to be overexpressed in *Xiphophorus* sword ^17^. In addition, other potassium channels, such as Kcnj13 and Kcnk5b, were also found to be responsible for zebrafish long fin mutants ^18–20^. Thus, these data suggest that KCNH genes, like other potassium channels, may be involved in embryonic developmental patterning and morphological diversity. However, the evolution of vertebrate KCNH genes and their systematic expression in zebrafish, a well-established vertebrate model system, are not well understood.

Here, we systematically cloned the zebrafish kcnh genes and analyzed vertebrate KCNH gene evolution using phylogenetic and synteny analyses. The three subfamilies (Kv10-12) were already separated before the origin of vertebrates, and each subfamily member was generated via whole-genome duplication (WGD) events during vertebrate evolution. We also examined their gene expression patterns during zebrafish embryogenesis and found that most are expressed in nervous tissues, consistent with their functions in neural physiology. In addition, *kcnh6a* and *kcnh7a* are expressed in the embryonic heart, and three genes (*kcnh2b*, *kcnh3*, and *kcnh6a*) were transiently expressed in non-neuronal somatic tissues, suggesting a unique function in embryonic development.

## RESULTS

### Cloning and validating zebrafish Kcnh channels

To identify zebrafish *kcnh* genes, we searched the NCBI and Ensembl databases (GRCz11) using BLAST with corresponding human *KCNH* genes as bait sequences. We identified 14 *kcnh* genes (**Table 1**), which were consistent with the current ZFIN (The Zebrafish Information Network). All 14 Kcnh channels share a conserved GFGN motif in the p-loop (**Fig. 1**). Since most of the channels were derived from bioinformatics analyses of databases, we confirmed the coding genes by RT-PCRs using a mixed cDNA pool from mixed-stage fish embryos (18-120 hours post fertilization). As typical protein and structural domains characterize the KCHN channels, we retrieved their protein structures from AlphaFold. As reported ^21^, all of the zebrafish Kcnh channels possess similar structures: PAS (Per-Arnt-Sim), N-linker, TM (transmembrane) domains(S1-S6), P-loop, c-linker, and CNBHD (cyclic nucleotide-binding homology domain) (**Fig. 2A-Q**). It is worth noting that the Kcnh6b lacks the PAS domain (**Fig. 2J**), consistent with a previous report ^22^.

**Figure 1.**
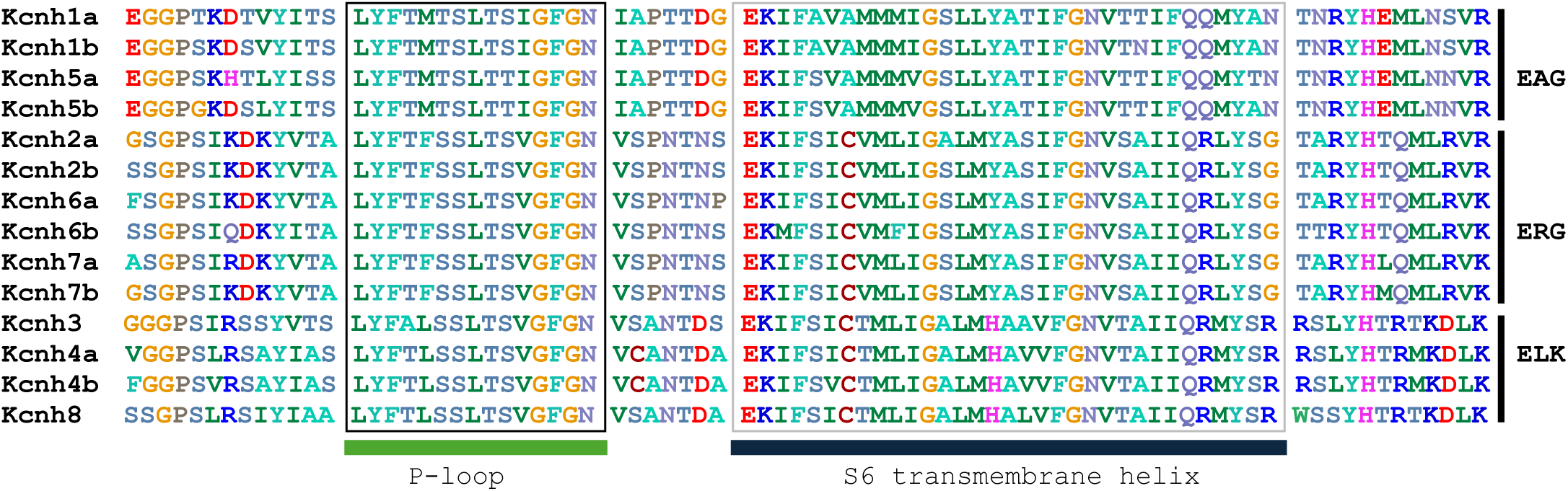
Alignment of zebrafish Kcnh channel P-loop and S4 domains. Zebrafish Kcnh channels possess one P-loop and six transmembrane domains. Only sequences around P-loop and S4 were shown. P-loop, the potassium pore domains (T-X-G-Y/F-G) were boxed in a black line. The S6 transmembrane domain next to the filter domain is also boxed in a gray line.

**Figure 2.**
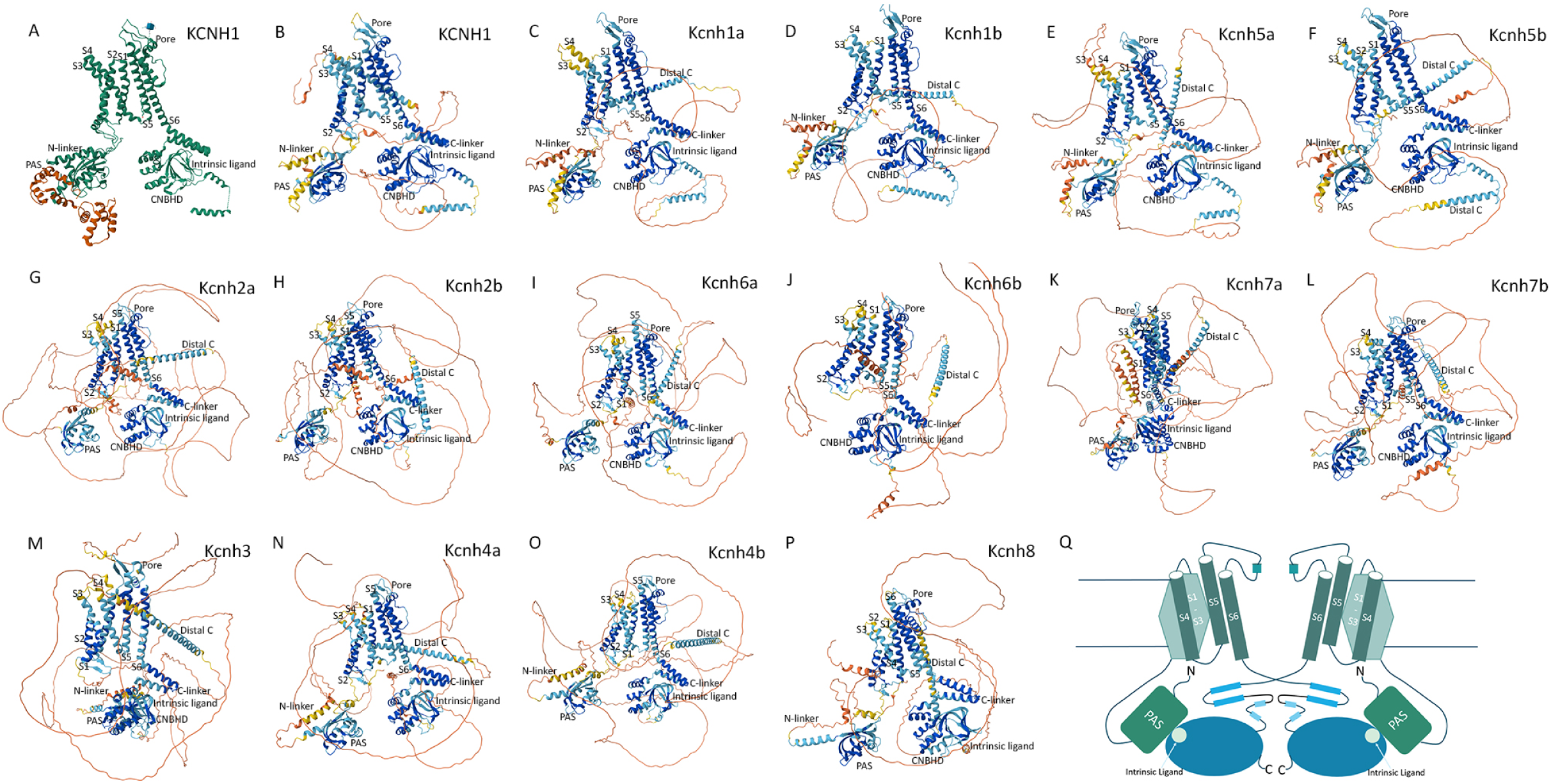
The zebrafish KCNH channels have protein structures similar to those of human KCNH1. **A.** Human KCNH1 subunit structure from an experiment in the RCSB PDB. **B.** KCNH1 structure predicted by AlphaFold. **C-P**. Predicted zebrafish Kcnh subunit protein structures. There is no PAS domain in zebrafish Kcnh6b. **Q.** Illustration of the KCNH tertiary structure, modified from the reference ^21^. The KCNH channel is a tetramer, but only two subunits are shown here. cNBHD (cyclic nucleotide-binding homology domain) is illustrated as a blue oval. A light blue circle indicates the intrinsic ligand-binding site. PAS, Per-ARNT-Sim domain. S1-S6, transmembrane domains.

**Table 1.**
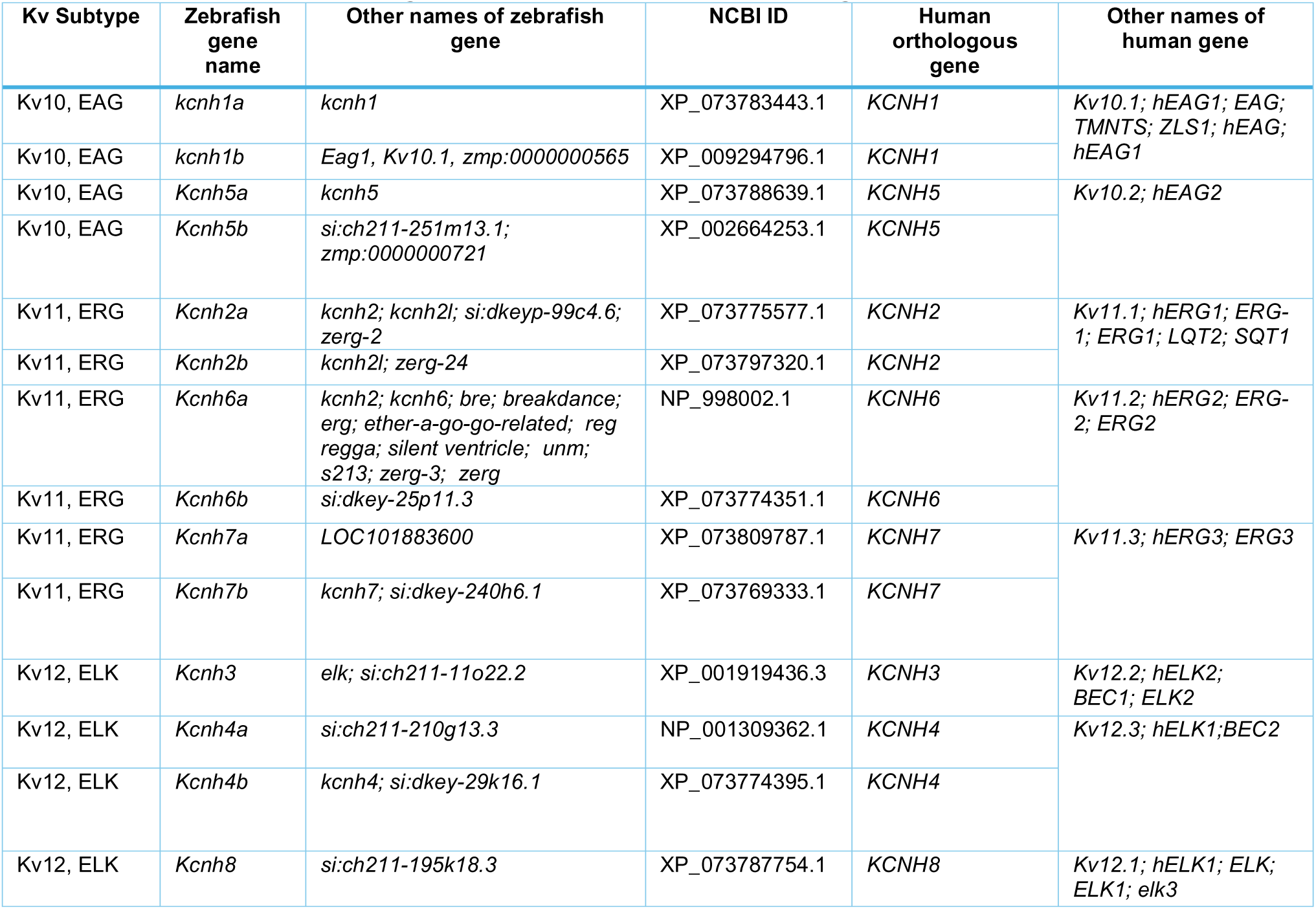
Zebrafish *kcnh* genes and their human orthologues.

### Evolutionary history of the Vertebrate KCNHs

There are usually more homologous genes in teleosts than in most tetrapods within a given gene family. Consistently, zebrafish have 14 *kcnh* genes, compared to 8 in humans (Table 1). This is a phenomenon mainly caused by the teleost-specific WGD that happened around 267-286 million years ago) ^23–27^. With the ohnologs (paralogues generated from WGD events), their gene names and relationships are usually confusing. To investigate the evolutionary origin of Kcnh channels and resolve their gene identities, we performed a phylogenetic analysis using KCNH channels from representative vertebrate species (**Table S1**). Consistent with the current classification, the KCNH channels were clustered into three distinct groups: Kv10 or EAG, Kv11 or ERG, and Kv12 or ELK (**Fig. 3**). Interestingly, invertebrate (fly, worm, lancelet, and tunicate) KCNH channels were clustered with their vertebrate orthologues. This suggests that the three subfamilies already evolved before the split of the vertebrate lineage, consistent with their origin back to the eumetazoan ^28^.

**Figure 3.**
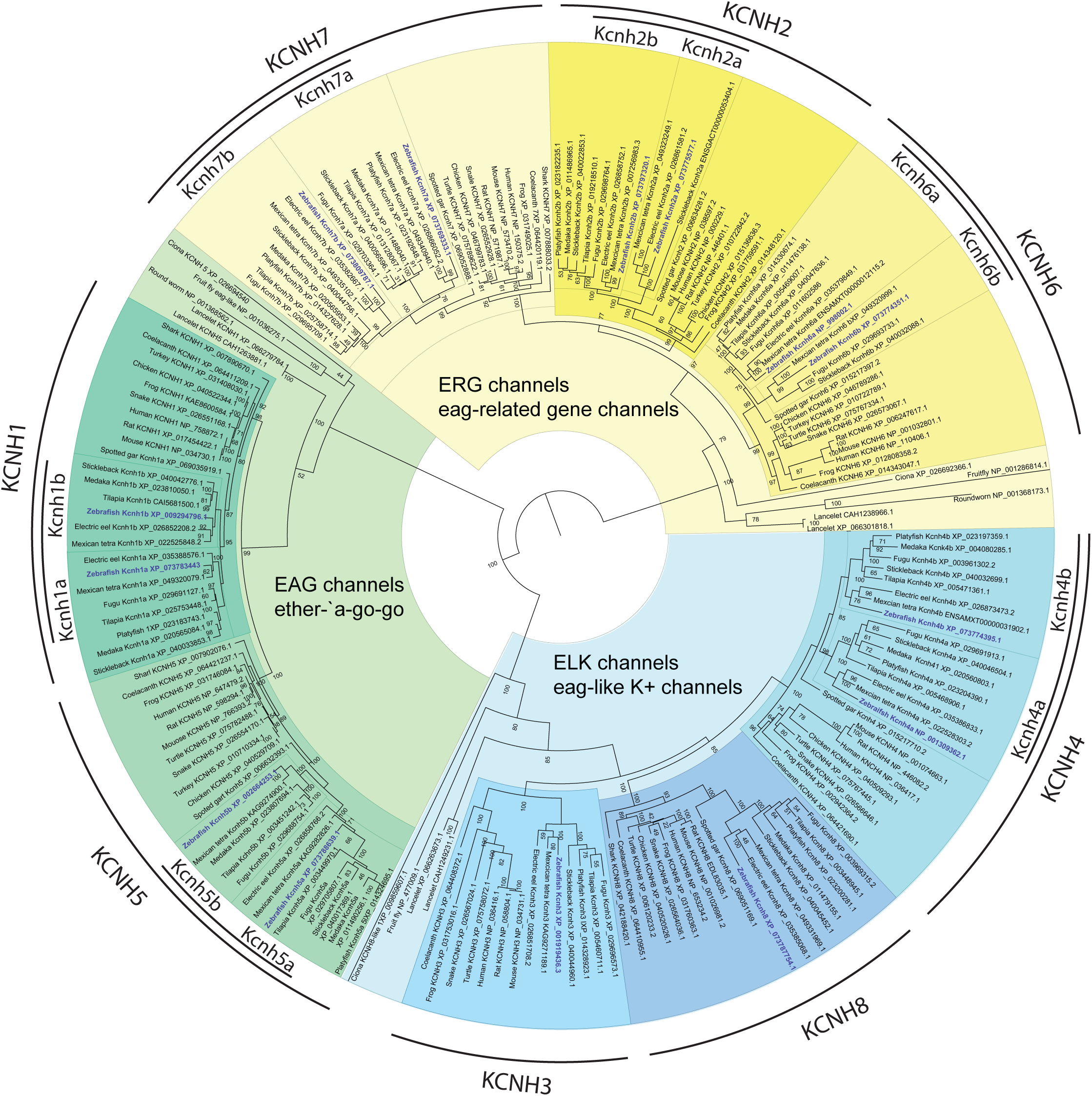
Maximum likelihood phylogeny of vertebrate Kcnh channels. Representative species (human, rat, mouse, chicken, turkey, frog, snake, turtle, coelacanth, zebrafish, tilapia, platy fish, stickleback, medaka, spotted gar, shark, lancelet, tunicate, roundworm, and fruit fly) were analyzed. Numbers at each node indicate the supporting values based on 1,000 bootstrap replicates. Branch lengths are proportional to the expected number of replacements per site. The phylogeny was inferred with the JTT model plus gamma distribution using RaxML(8.2.12). The three subtypes of Kcnh channels were grouped together and indicated with different colored shades. The zebrafish Kcnh channels were highlighted in blue. The channel protein name is labeled on the edge of the phylogenetic tree. The protein sequence reference number is included in the phylogenetic tree to avoid confusion when genes are duplicated. The duplicated channels are indicated by black lines next to their names, ending with a and b.

### The KCNH channels were duplicated in teleosts

The EAG subgroup comprises KCNH1 and KCNH5 channels. Both KCNH1 and KCNH5 channels formed a distinct subgroup within this cluster, with the invertebrate KCNH1/5-like channels serving as the outgroup (Fig. 3, green shading). There was a single KCNH1 and KCNH5 channel in tetrapods. In contrast, two copies of these Kcnh1 and Kcnh5 channels were present in multiple teleost species, and they were clustered closer than their tetrapod orthologues. Thus, our phylogenetic analysis suggested that these were ohnologues from teleost-specific WGD. To further test the relationships between tetrapod and teleost EAG channels, we then checked the syntenies of the KCNH1 and KCNH5 genes in a few representative vertebrate species (**Fig. 4A-B**). The *KCNH1* gene is in a conserved synteny, *SERTAD4-HHAT-KCNH1-RCOR3-TRAF5*, which shark, mouse, and human share (**Fig. 4A**). In spotted gar, *SERTAD4* is replaced by *pex6*. In comparison, this synteny is broken into two pieces in zebrafish: *kcnh1a-hhat* and *kcnh1b-pex6*, suggesting that the gene locations were reshuffled after the teleost-specific WGD. Similarly, the *KCNH5* gene is also within a conserved synteny: *PRKCH-HIF1A-SNAPC1-SYT16-KCNH5-RHOJ-GPHB1-PPP2R5E-WDR89* (shark, spotted gar, mouse, and human). In zebrafish, there are two smaller ones: *kcnh5a-rhoj-ppp2r5ea-snapc1a-hif1ab-prkchb* and *gphb5-kcnh5b-ppp2r5eb-wdr89* (**Fig. 4b**). The shared content, differential truncation, and reshuffling of these syntenies supported the idea that the duplicated *kcnh1* and *kcnh5* were generated through the teleost-specific WGD.

**Figure 4.**
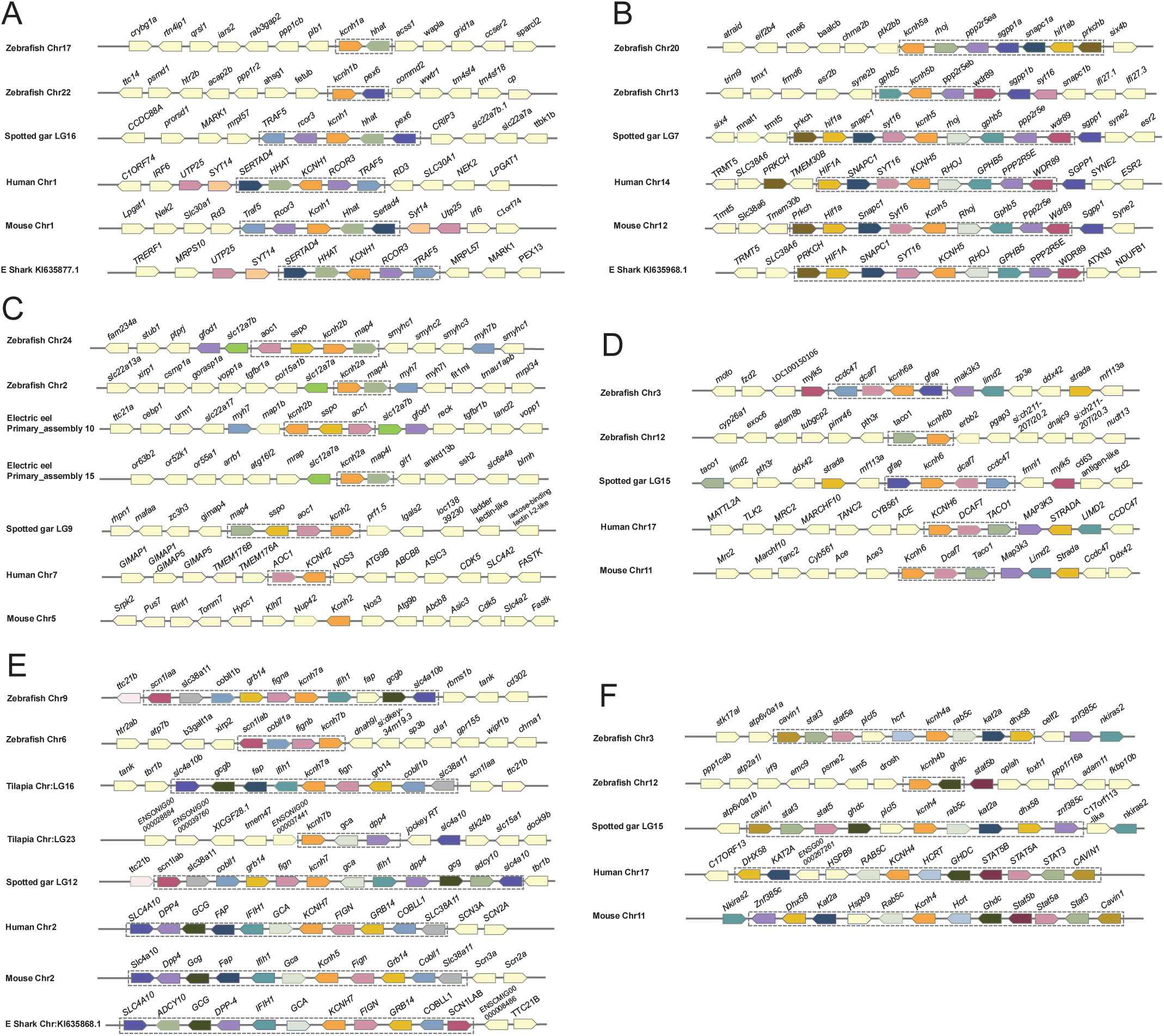
The syntenic analyses of the duplicated *kcnh* genes in representative vertebrate species. **A.** Vertebrate *kcnh2* synteny: *traf5-hhat-kcnh1-rcor3-hhat*. The zebrafish *kcnh1a* gene is linked to hhat, as in sharks and humans. **B.** Vertebrate *kcnh5* synteny is relatively stable in shark, spotted gar, human, and mouse. In zebrafish, this conserved synteny is split into two short ones with *kcnh5a* and *kcnh5b*. **C.** the *kcnh2* synteny, *map4-sspo-aoc1-kcnh2* is kept well in zebrafish and electric eel compared to the spotted gar. Only *map4l* is linked with *kcnh2b* in both zebrafish and electric eel. The human *KCNH2* is linked only to AOC1, whereas the mouse Kcnh2 is not linked to any other genes in the spotted gar synteny. **D.** Vertebrate *kcnh6* synteny, *kcnh6-dcaf7-taco1*. In zebrafish, *kcnh6a* is linked to *dcaf7,* and *kcnh6b* is linked to *taco1*. **E.** Vertebrate *kcnh7* synteny is relatively long and has 11 genes in the elephant shark and spotted gar genomes. The *kcnh7a* is linked to the majority member of the synteny, while the *kcnh7b* is linked to two or three genes in both zebrafish and tilapia. **F.** Vertebrate *kcnh4* synteny is also relatively conserved. The zebrafish *kcnh4a* gene is located within a longer syntenic block than the *kcnh4b* gene. The duplicated *kcnh* genes were highlighted in orange. The illustration of the genes and their sizes is not proportional to the distances between neighboring genes. Due to limited genome annotation, we do not have access to syntenies in the elephant shark, which is usually the primary condition in **C**, **D**, and **F**.

In the ERG subgroup, there are 3 members: KCNH2, KCNH6, and KCNH7. All vertebrate EAG channels formed an independent clade, with invertebrate EAG channels as the outgroup (Fig. 3, yellow shading). Interestingly, most teleost species had two copies of these three channels, forming a distinct clade compared to tetrapod channels. Thus, they were likely orthologs from the teleost-specific WGD. Next, we performed syntenic analysis. In the shark genome, we did not detect the KCNH2 and KCNH6 genes, likely due to incomplete annotation. However, we did find shared syntenic content among spotted gar (*map4-sspo-aoc1-kcnh2*), zebrafish (*aoc1-sspo-kcnh2b-map4*, *kcnh2a-map4l*), and electric eel (*kcnh2b-sspo-aoc1*, *kcnh2a-map4l*) (**Fig. 4C**). The *KCNH6* gene is in a conserved synteny block represented by spotted gar, *gfap-kcnh6-dcaf7-ccdc47*. In zebrafish, *kcnh6a* is in the synteny, whereas *kcnh6b* is not, suggesting shuffling of gene positions (**Fig. 4D**). The synteny of KCNH7 (*slc4a10-adcy10-gcg-dpp4-ifih1-cga-kcnh7-fign-grb14-cobll1-scn1lab*) is shared among humans, mice, spotted gars, and sharks (**Fig. 4E**). In teleost (tilapia and zebrafish), the *kcnh7a* gene is in a less interrupted shared synteny. In contrast, the *kcnh7b* gene is in a smaller, fragmented syntenic region (zebrafish: *kcnh7b-fign-cobll1a-scn1lab*; tilapia: *kcnh7b-gca-dpp4*). These shared syntenies suggest that the 3 EAG duplicates were generated from the teleost ancestor. It is worth noting that *kcnh7b* was named before *kcnh7a*, which is absent in GRCz11, but was recently identified as "*kcnh7b*" in GRCz12 ^29^. Here, we named *kcnh7a* and *kcnh7b* based on the integrity and size of shared synteny ^30^.

The ELK channel subgroup has three members: KCNH3, KCNH4, and KCNH8. All the vertebrate ELK channels were clustered with the invertebrate ELK channels, parallel to the EAG and ERG channels (**Fig. 3**, blue shade). KCNH3, KCNH4, and KCNH8 formed a distinct clade from each other. All the representative vertebrate species have one copy of KCNH3 and KCNH8. In contrast, most teleosts have two copies of the Kcnh4 channel, whereas tetrapods have only one. To further test whether they were from the teleost WGD, we turned to synteny analysis. Although information on chondrichthyans is limited, we identified a shared synteny among humans, mice, and spotted gars: *cavin1-stat3-stat5-ghdc-plcl5-kcnh4-rab5c-kat2-dhx58-znrf384c* (**Fig. 4F**). In zebrafish, *kcnh4a* remains in this synteny, whereas kcnh4b is linked only to *ghdc*, suggesting that they are ohnologues from the teleost WGD. It is worth noting that the *STAT5* gene was duplicated in humans and mice (*STAT5A* and *STAT5B*), likely through tandem duplication, as the two duplicates are located next to each other within this synteny.

### The gene expression of KCNH channels during zebrafish embryogenesis

To further understand the relationships among the three subtypes of KCNH Kv channels, we examined gene expression in zebrafish embryos by whole-mount *in situ* hybridization, as spatial and temporal expression patterns often indicate functions. The *kcnh1a* gene was detected in the notochord and hindbrain of 15-somite-stage (15S) embryos (**Fig. 5A, E**). Its expression receded and was limited to the posterior of the notochord at 24 hpf (hours post fertilization). It also became evident in the central nervous system. Particularly, it was expressed in the Rohon-Beard neurons of the neural tube (**Fig. 5B, F**). Upon 48 hpf, the notochord and neural tube expression withdrew, and the brain expression remained at both 48 hpf and 72 hpf. Meanwhile, this gene was also found in the olfactory organs and trigeminal ganglia (**Fig. 5C-D, G-H**). Consistent with the former report ^31^, *kcnh1a* was also detected in the epiphysis (pineal gland) from 24-72dpf (**Fig. 5C-D, G-H**). The *kcnh1b* gene was expressed weakly in whole embryos at 15S (**Fig. 5I, M**). It became restricted to the brain and the dorsal-lateral neural tube, mostly in the Rohon-Beard neurons, but not in the notochord, at 24hpf (**Fig. 5J, N**). Brain expression remained strong from 48 to 72 hpf. Still, the neural tube expression retracted from the spinal cord at 72 hpf (**Fig. 5K-L, O-P**). The neural expression of *kcnh1a* and *kcnh1b* is consistent with their functions in neurogenesis^31^. The *kcnh5a* gene was expressed in the anterior parts of the hindbrain and spinal cord at 15S (**Fig. 5Q, U**), and its expression was temporarily expanded to the dorsal neural tube, likely including the Rohon-Beard neurons, at 24 hpf (**Fig. 5R, V**). Then, this neural tube expression domain retracted at 48 hpf, in contrast to the evident expression in the head region (**Fig. 5S, W**). At the stage of 72 hpf, the *kcnh5a* was also detected in the retina (**Fig. 5T, X**). The *kcnh5b* gene was not detectable at 15S, but weakly found in the base of the brain at 24 hpf (**Fig. 5Y-Z, CC-DD**). At 48 hpf, *kcnh5b* was detected in the olfactory placodes and facial sensory ganglion (**Fig. 5AA, EE**). This expression domain persisted at 72 hpf, and new domains, such as the retina and the midbrain optic tectum, became evident (**Fig. 5BB, FF**).

**Figure 5.**
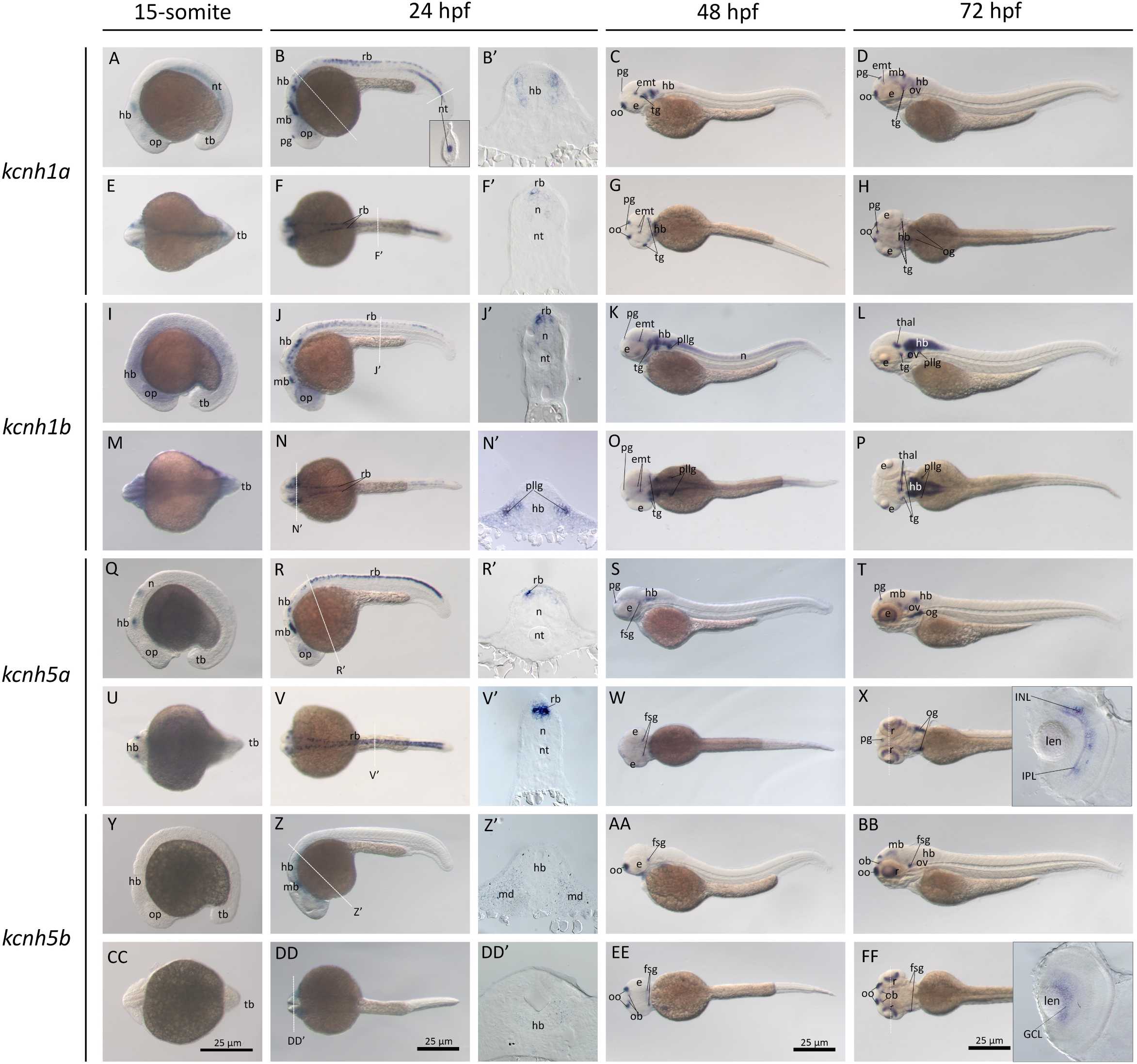
The *kcnh1a*, *kcnh1b*, *kcnh5a*, and *kcnh5b* gene (EAG channel) expression during zebrafish embryogenesis. Whole-mount *in situ* hybridization of zebrafish embryos at 15 somite stage, 15S (**A, E, I, M, Q, U, Y, CC**), 24 hpf (**B, F, J, N, R, V, Z, DD**), 48 hpf (**C, G, K, O, S, W, AA, EE**), and 72 hpf (**D, H, L, P, T, X, BB, FF**). The anterior is to the left in all the whole-mount images, and the dorsal is to the top in all transverse sections (**B’, F’, G’, J’, N’, O’, R’, V’, W’, Z’, DD’**). The white dashed lines indicate the positions of sections left to the panel or transverse sections in the inserts of corresponding panels. The letters below or around the dashed lines correspond to the section panels. **A-D**. Lateral view of gene expression of *kcnh1a*. **E-H**. Dorsal view of gene expression of *kcnh1a*. **I-L**. Lateral view of gene expression of *kcnh1b*. **M-P**. Dorsal view of gene expression of *kcnh1b*. **Q-T**. Lateral view of gene expression of *kcnh5a*. **U-X**. Dorsal view of gene expression of *kcnh5a*. **Y-BB**. Lateral view of gene expression of *kcnh5b*. **CC-FF**. Dorsal view of gene expression of *kcnh5b*. Scale bars are at the bottom row of images. 250Lμm for whole-mount images. *e*, eye; *emt*, eminentia thalami; *fsg*, facial sensory ganglion; *GCL*, ganglion cell layer; *hb*, hindbrain; *INL*, inner nuclear layer; *IPL*, Inner plexiform layer; *len*, lens; *mb*, midbrain; *mc*, mesenchymal; *n*, neural tube; *nt*, notochord; *ob*, olfactory bulbs; *ofp*, olfactory placode; *og*, octaval ganglion; *op*, optic vesicles; *ov*, otic vesicles; *pg*, pineal gland; *pllg*, posterior lateral line ganglia; *r*, retina; *rb*; Rohon-Beard neurons; *tb*, tail bud; *tg*, trigeminal ganglion; *thal*, thalamus.

ERG gene expression in zebrafish embryos is relatively diverse since there are 6 genes in this subgroup. The *kcnh2a* gene was weakly detected in the head region of the 15S stage fish embryos, but it became clear in the brain and spinal cord at 24hpf (**Fig. 6A-B, E-F**). Its expression further narrowed to some neural ganglia and the thymus, in addition to the neural tube, at 48 hpf (**Fig. 6C, G**). At 72 hpf, neural ganglia remained (**Fig. 6D, H**). The *kcnh2b* gene was mainly found in the neural tube at 15S (**Fig. 6I, M**). This gene was then expressed in the forebrain, hindbrain, and Rohon-Beard neurons in the neural tube at 24hpf (**Fig. 6J, N**). It was then limited to specific cranial ganglia in the brain at 48 and 72 hpf (**Fig. 6K-L, O-P**). The *kcnh6a* gene was strongly expressed in somites at 15S (**Fig. 6Q, U**), and was limited to the dermomyotome at 24 hpf (**Fig. 6R, V**). In addition, it was also found in the primitive heart tube at this stage (**Fig. 6R**). At 48 hpf, *kcnh6a* was limited to the heart and forebrain (**Fig. 6S, W**). This forebrain expression was transient and retracted at 72hpf, and the gene was detected only in the heart (**Fig. 6T, X**). The *kcnh6b* gene was barely detected in the whole fish embryos at 15S (**Fig. 6Y, CC**), but was evident in the midbrain and hindbrain at 24hpf (**Fig. 6Z, DD**). The broad head expression remains in the whole brain from 48-72hpf (**Fig. 6AA-BB, EE-FF**). The *kcnh7a* gene was not detectable at 15S (**Fig. 6GG, KK**), but was found evident in the head at 24hpf (**Fig. 6HH, LL**). Then this gene maintained its brain expression from 48 to 72 hpf (**Fig. 6II-JJ, MM-NN**). The *kcnh7b* gene was expressed weakly in the hindbrain at 15S (**Fig. 6OO, SS**). Then, at 24hpf, it was found throughout the central nervous system, including Rohon-Beard neurons, in addition to the presomatic mesoderm (**Fig. 6PP, TT**). The neural tube expression retracted at 48hpf, but its brain expression remained strong from 48 to 72 hpf (**Fig. 6QQ-RR, UU-VV**).

**Figure 6.**
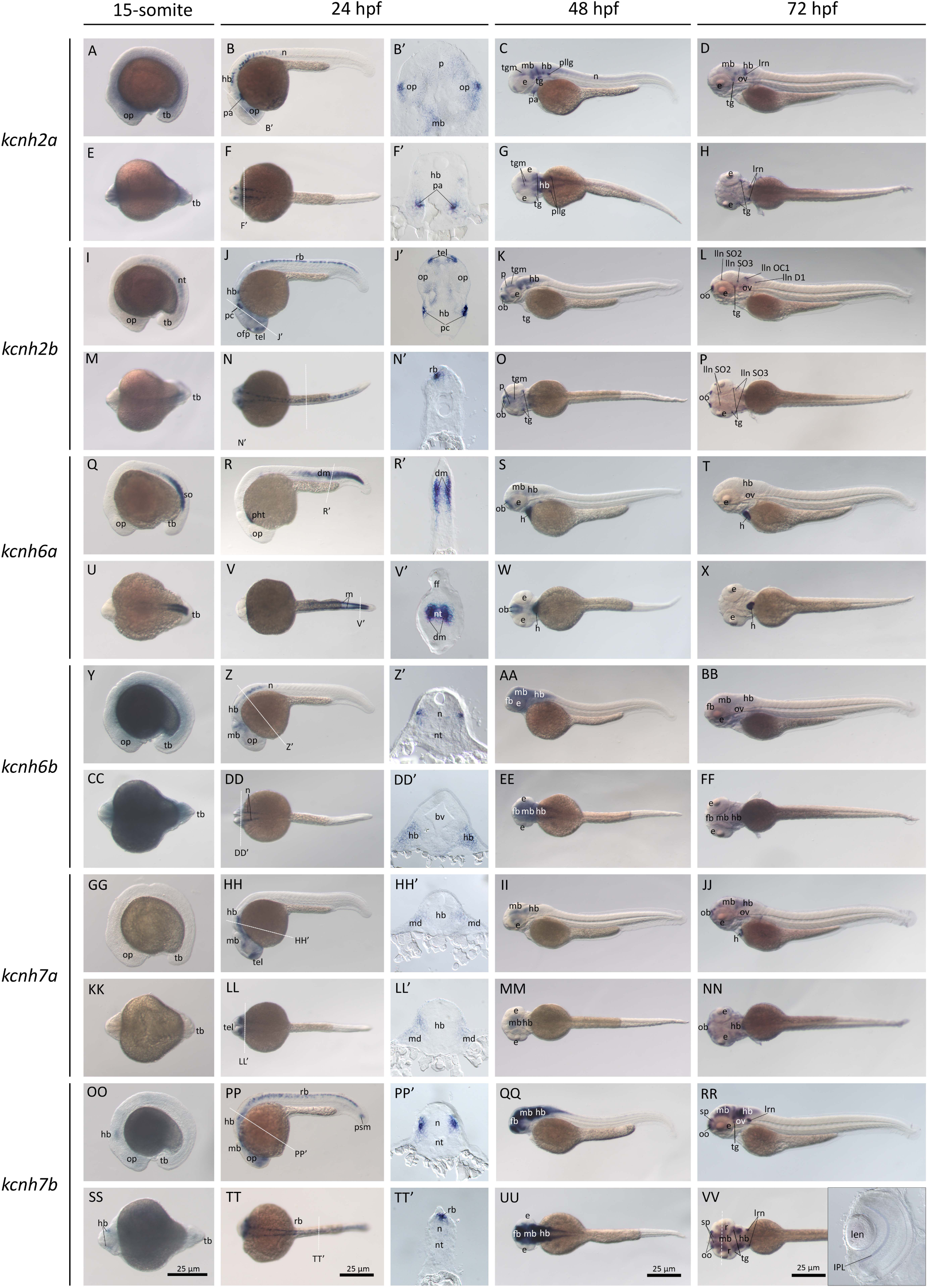
Gene expression of the *kcnh2a*, *kcnh2b*, *kcnh6a*, *kcnh6b, kcnh7a,* and *kcnh7b* genes (ERG channel) during zebrafish embryogenesis. Whole-mount *in situ* hybridization of zebrafish embryos at 15S stage (**A, E, I, M, Q, U, Y, CC, GG, KK, OO, SS**), 24 hpf (**B, F, J, N, R, V, Z, DD, HH, LL, PP, TT**), 48 hpf (**C, G, K, O, S, W, AA, EE, II, MM, QQ, UU**), and 72 hpf (**D, H, L, P, T, X, BB, FF, JJ, NN, RR, VV**). The anterior is to the left in all the whole-mount images, and the dorsal is to the top in all transverse sections (**B’, F’, G’, J’, N’, O’, R’, V’, W’, Z’, DD’, HH’, LL’, PP’, TT’**). The white dashed lines indicate the positions of the sections to the left of the panel. The letters below or around the dashed lines correspond to the section panels. **A-D**. Lateral view of gene expression of *kcnh2a*. **E-H**. Dorsal view of gene expression of *kcnh2a*. **I-L**. Lateral view of gene expression of *kcnh2b*. **M-P**. Dorsal view of gene expression of *kcnh2b*. **Q-T**. Lateral view of gene expression of *kcnh6a*. **U-X**. Dorsal view of gene expression of *kcnh6a*. **Y-BB**. Lateral view of gene expression of *kcnh6b*. **CC-FF**. Dorsal view of gene expression of *kcnh6b*. **GG-JJ**. Lateral view of gene expression of *kcnh7a*. **KK-NN**. Dorsal view of gene expression of *kcnh7a*. **OO-RR**. Lateral view of gene expression of *kcnh7b*. **SS-VV**. Dorsal view of gene expression of *kcnh7b*. Scale bars are at the bottom row of images. 250Lμm for whole-mount images. *bv*, brain vesicles*; dm*, dermomyotome; *e*, eye; *emt*, eminentia thalami; *fb*, forebrain; *fsg*, facial sensory ganglion; *h*, heart; *hb*, hindbrain; *lln*, lateral line neuromast (*d*, dorsal*; io*, infraorbital; *oc*, Occipital, *so*,supraorbital); *len*, lens; *lrn*, lateral reticular nucleus; *mb*, midbrain; mc, mesenchymal; *n*, neural tube; *nt*, notochord; *ob*, olfactory bulbs; *ofp*, olfactory placode; *og*, octaval ganglion; *op*, optic vesicles; *ov*, otic vesicles; *p*, pallium; *pa*, pharyngeal arches; *pc*, placede; *pg*, pineal gland; *pht*, primitive heart tube; *pllg*, posterior lateral line ganglia; *r*, retina; *rb*; Rohon-Beard neurons; *so*, somite; *tb*, tail bud; *tg*, trigeminal ganglion; *tgm*, tegmentum; *tel*, telencephalon; *thal*, thalamus.

Among the ELK genes, *kcnh3* was expressed in the hindbrain and anterior of the neural tube at 15S (**Fig. 7A, E**). These expression domains were retained at 24 hpf, while the gene was also found at the base of the forebrain and was notably expressed in the anterior dermomyotome (**Fig. 7B, F**). Upon entering the 48 hpf stage, the gene was found dominantly in the central nervous system, including the neural tube (**Fig. 7C, G**). At 72 hpf, *kcnh3* expression was decreased and was limited to specific regions of the brain, such as the olfactory vesicles and bulbs, and the pineal gland (**Fig. 7D, H**). The *kcnh4a* gene was expressed in the brain at 15S and persisted until 24 hpf (**Fig. 7I-J, M-N**). In addition, it was weakly detected in some Rohon-Beard neurons in the anterior region of the neural tube (**Fig. 7J, N**). At 48hpf, *kcnh4a* was expressed at the olfactory bulbs and HCRT neurons between eyes (**Fig. 7K, O**), as reported to be related to sleep modulation in zebrafish^32^. These domains remained at 72 hpf (**Fig. 7L, P**). The *kcnh4b* gene was also weakly expressed in the brain and notochord at 15S (**Fig. 7Q, U**). The brain showed increased expression, and neural tube expression decreased. At 48hpf, *kcnh4b* was limited to olfactory placodes and the pineal gland (**Fig. 7S, W**). These expression domains persisted at 72 hpf, and *kcnh4b* mRNA began to be expressed in the midbrain (**Fig. 7T, X**). The *kcnh8* gene was not detectable at 15S (**Fig. 7Y, CC**), but was found at the base of the brain at 24 hpf (**Fig. 7Z, DD**). At 48hpf, *kcnh8* was expressed at the pallium, lateral of the midbrain, and the anterior of the hindbrain (**Fig. 7AA, CC**). In contrast, it was limited to the anterior part of the hindbrain at 72hpf (**Fig. 7BB, DD**).

**Figure 7.**
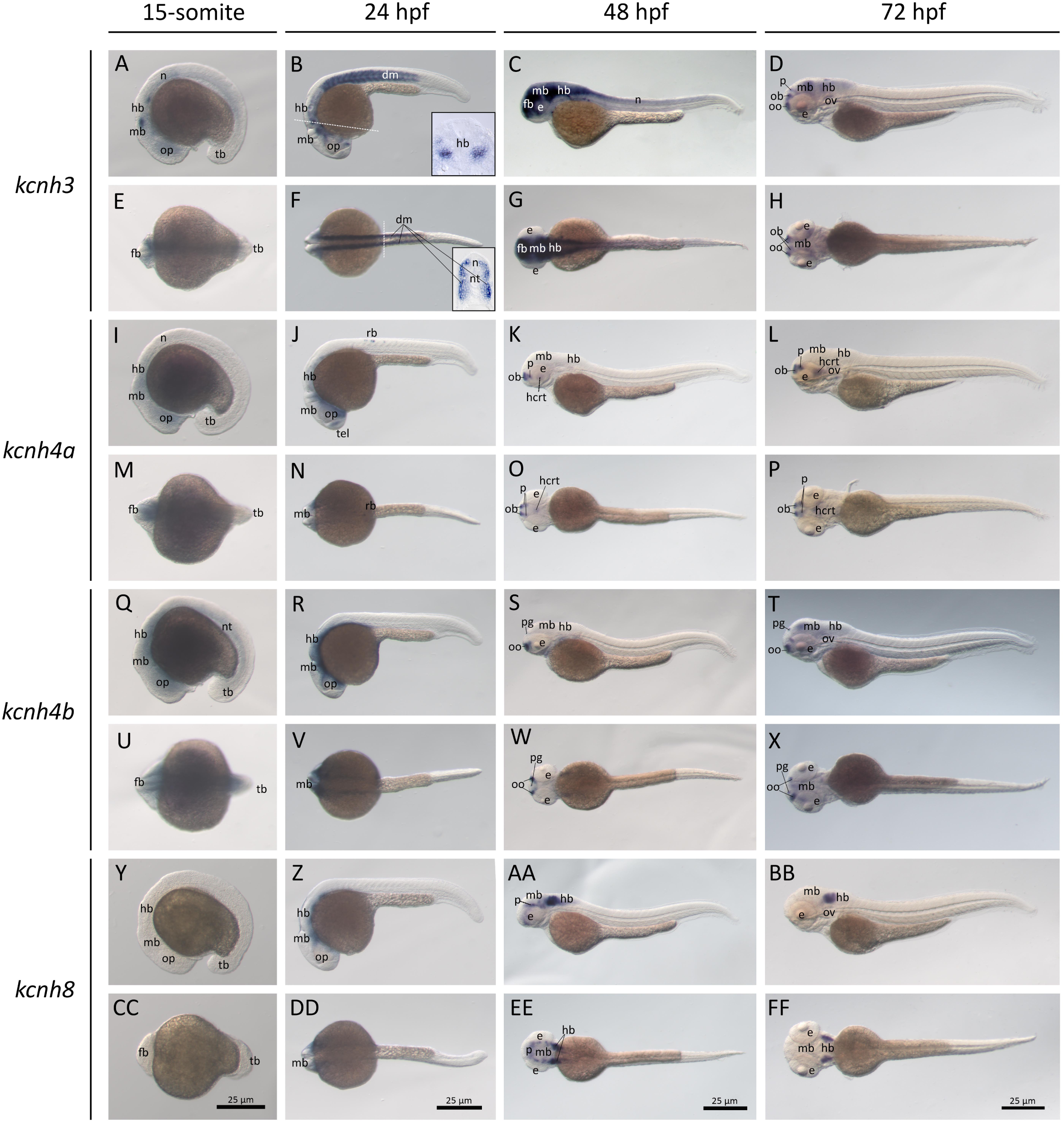
The gene expression of the ELK channel genes, *kcnh3*, *kcnh4a*, *kcnh4b*, and *kcnh8,* during zebrafish embryogenesis. Whole-mount *in situ* hybridization of zebrafish embryos at the 5S stage. (**A, E, I, M, Q, U, Y, CC**), 24 hpf (**B, F, J, N, R, V, Z, DD**), 48 hpf (**C, G, K, O, S, W, AA, EE**), and 72 hpf (**D, H, L, P, T, X, BB, FF**). The anterior is to the left in all the whole-mount images. The white dashed lines indicate the positions of the transverse sections in the inserts of corresponding panels (**B, F**). **A-D**. Lateral view of gene expression of *kcnh3*. **E-H**. Dorsal view of gene expression of *kcnh3*. **I-L**. Lateral view of gene expression of *kcnh4a*. **M-P**. Dorsal view of gene expression of *kcnh4a*. **Q-T**. Lateral view of gene expression of *kcnh4b*. **U-X**. Dorsal view of gene expression of *kcnh4b*. **Y-BB**. Lateral view of gene expression of *kcnh8*. **CC-FF**. Dorsal view of gene expression of *kcnh8*. Scale bars are at the bottom row of images. Scale bars are at the bottom row of images. 250Lμm for whole-mount images. *e*, eye; *fb*, forebrain, *hb*, hindbrain; *hcrt*, hypocretin (HCRT) neurons in the hypothalamus, *mb*, midbrain; *n*, neural tube; *nt*, notochord; *ob*, olfactory bulbs; *ofp*, olfactory placode; *op*, optic vesicles; *ov*, otic vesicles; *p*, pallium; *pg*, pineal gland; *tb*, tail bud; *tel*, telencephalon.

## DISCUSSION

The KCNH voltage-gated potassium channels play essential roles in repolarization during an action potential. Thus, this group of potassium channels has been frequently reported in human neural and cardiovascular physiology and disease. Here, we systematically examined 14 Kcnh channel gene expression in zebrafish embryos and analyzed the evolutionary history of vertebrate Kcnh channels.

The KCNH channels have a deep origin that can be traced back to metazoan ^28^. The EAG, ERG, and ELK subgroups were already established before the split of vertebrates. However, the number of invertebrate KCNH channels is much less than that of vertebrates. Compared with tetrapods, teleosts have more Kcnh channels. This phenomenon is consistent with many gene families, including Hox genes and other potassium channel genes ^33–37^. It is well known that the increase in gene numbers has been associated with the two successive rounds of WGDs (1R and 2R) in the vertebrate ancestor about 500 million years ago^38–40^. There is only one Hox gene cluster in most invertebrates, such as the fruit fly, tunicate, and amphioxus. In contrast, 4 Hox gene clusters are present in the mouse and human genomes ^41^. The current consensus is that a few more WGD events occurred within vertebrates ^42^. Notably, the ancestor of teleost underwent another WGD (3R), leading to an increase in gene number within this lineage around 320 million years ago ^27,43^. After WGDs, ohnologs could be retained or lost in each species. Thus, the number of paralogous genes may vary across lineages ^25,44^. Our phylogenetic results are consistent with this notion, and we proposed a model for the evolution of the vertebrate KCNH channel (**Fig. 8A-B**). In most chordates, there is only one gene for EAG, ERG, and ELK. After the two rounds of WGDs, the number of each channel increased to four. However, only one KCNH1 and KCNH5 were retained, and the other ohnologs were lost. Another scenario is that four ohnologs were generated by the WGDs, but one was lost. So only three ohnologs in ERG (KCNH2, KCNH6, and KCNH7) and ELK (KCNH3, KCNH4, and KCNH7). These KCNH channels were further duplicated in teleosts by the 3R WGD after the teleost split. Immediately, gene loss occurred only in the ELK subfamily, leading to two copies of Kcnh4a and Kcnh4b. However, there is only one Kcnh3 and one Kcnh8 (**Fig. 3**). Consistently, our syntenic analyses also support this model, as the duplicated genes are located in evolutionarily conserved syntenies in the analyzed teleost species.

**Figure 8.**
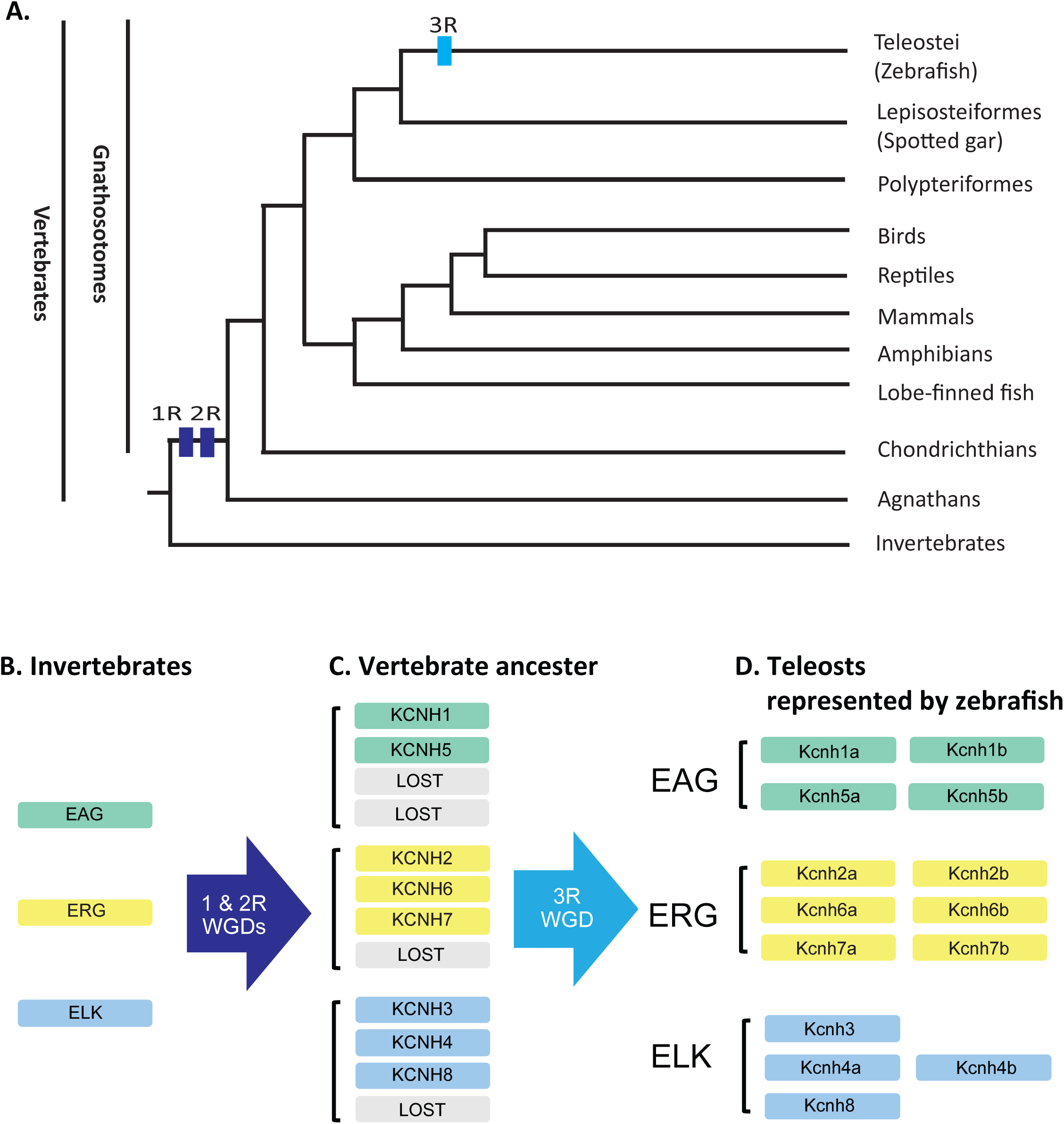
Model of vertebrate *Kcnh* gene evolution. Based on our phylogenetic and syntenic analysis, we proposed an evolutionary model for vertebrate potassium channels. **A.** Whole genome duplication events (1R-3R) in vertebrates. The horizontal blue bars indicate the WGD events. The vertebrate phylogenetic tree is illustrated based on the evolution of fibrillar collagens ^53,54^. **B.** The three Kv subtypes (EAG, ERG, and ELK) have already evolved in invertebrates. There is at least one gene for each subtype in most invertebrates. **C.** Two consecutive WGDs (R1 and R2) happened at the dawn of vertebrates. So, all the *Kcnh* genes were doubled twice in the vertebrate ancestor. Some of the duplicated genes were lost after the two WGDs: 2 in the EAG subfamily, 1 in the ERT subfamily, and 1 in the ELK subfamily. **D.** The teleost-specific WGD further reshaped the gene numbers in this group of animals. Most gene duplicates were maintained, except that one copy of kcnh3 and one copy of *kcnh8* were lost in teleosts.

Although further work is required to validate and refine the detailed expression profiles of these *kcnh* genes within the specified embryonic structures, the catalog presented here provides an essential foundation for such inquiry. Overall, we observed overlapping and distinct gene expression domains of the zebrafish duplicated *kcnh* channel genes in the nervous system (**Fig. 9**), suggesting that their primary functions could be complementary to K2P channels in neural excitability ^33^. It is worth noting that a few of them (*kcnh1a*, *kcnh1b*, *kcnh2b*, *kcnh4a*, *kcnh5a*, and *kcnh7b*) are expressed in Rohon-Beard neurons, the transient and the earliest mechanosensory neurons that are essential for the establishment of the rudimentary motor response circuit ^45^. This indicates that these Kcnh channels may play important roles in neurogenesis. Also, a few Kcnh channels (*kcnh1a, kcnh5b, kcnh2b, kcnh4a*, and kcnh4b) were found in the olfactory organs and bulbs, suggesting a role in the olfactory system that has not yet been reported. Similarly, three of them (*kcnh5a, kcnh5b*, and *kcnh7b*) were expressed in distinct retinal layers and play functional roles in zebrafish eye development and physiology.

**Figure 9.**
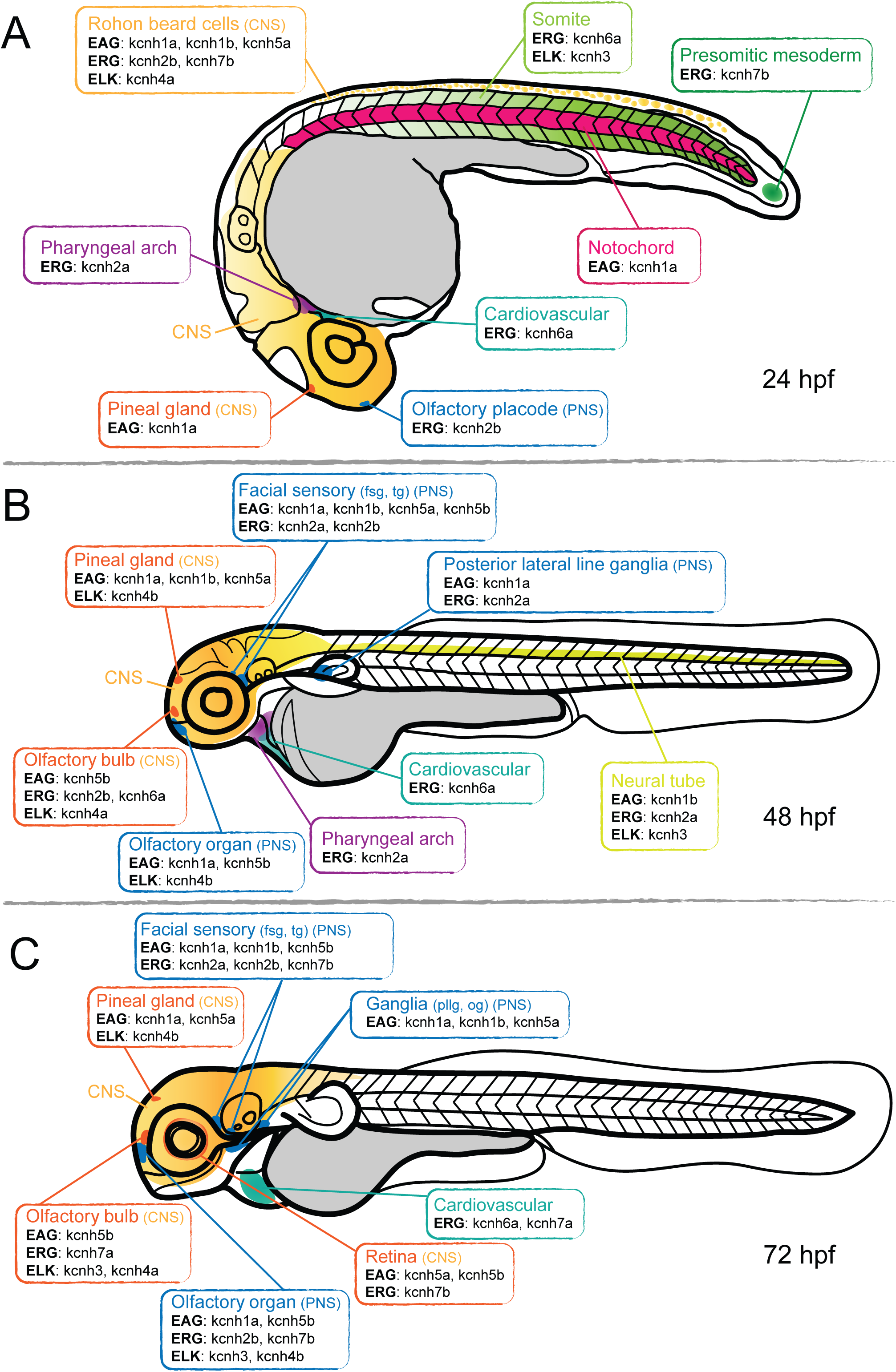
Schematic summary of the *kcnh* gene expression in 24-72 hpf zebrafish embryos. Only major embryonic structures are illustrated. The representative embryonic organs or tissues are color-coded. The *kcnh* genes expressed in the corresponding regions are listed in boxes of the same color. **A.** 24hpf. **B.** 48hpf. **C.** 72hpf. The size of the illustration is not proportional to the actual zebrafish embryos.

One common feature of the 3 subtypes of *kcnh* gene expression in zebrafish embryos is that duplicated genes share the same expression domains but also have unique domains. For example, *kcnh1a* and *kcnh1b* are expressed in the Rohon-Beard neurons of the neural tube. In contrast, only *kcnh1a* is expressed in the olfactory vesicles, and *kcnh1b* has a broader expression domain in the hindbrain at 72hpf. Another example is the *kcnh7a* and *kcnh7b*. Both are expressed in the brain, while only *kcnh7b* is present in the neural tube. This phenomenon could be explained by subfunctionalization. Based on the DDC (duplication-degeneration-complementation) model, both regulatory and protein-coding regions are duplicated by WGDs ^46^. The coding region evolves more slowly than the regulatory region due to constraints on protein function. Eventually, the splitting of the regularly expressed region leads to subfunctionalization. The subfunctionalization not only helps us understand gene functional redundancy but may also provide a unique opportunity to decipher how the gene functions of its mammalian orthologue are developmentally lethal. Another scenario is neofunctionalization, in which a duplicate acquires a new regulatory module, creating new gene expression domains. KCNH2 and KCNH6 could be the case after the two vertebrate WGDs, since KCNH2 is found in the mammalian heart, while Kcnh6a is the only KCNH in the zebrafish heart, as reported previously ^22,47^. However, it is hard to tell which one is new, since there is no reference in outgroup species such as sharks and other basal vertebrates.

Among the 14 kcnh genes, a few are also expressed in nonneural tissues. These nonneural tissues suggest they may have essential functions in embryonic differentiation and patterning. The most noticeable ones are the *kcnh6a* and *kcnh7a*, which are unique in that they are mainly expressed in the heart. Kcnh6a is consistent with previously reported findings, while Kcnh7a is a new channel in the zebrafish heart ^22,47^. Thus, it may lead to cautious use of zebrafish to model human long QT syndrome, since KCNH2 is the functional counterpart in human hearts. In addition, *kcnh2a* was expressed in the pharyngeal arch, and *kcnh7b* in the presomatic mesoderm, suggesting their roles in embryonic development and physiology. We also observed that *kcnh1a, kcnh2b,* and *kcnh4b* in the notochord and *kcnh3* and *kcnh6a* in the dermomyotome, a somite compartment that is a key regulatory domain for zebrafish fin size. We have found that transient ectopic *kcnj13* expression in somites induced elongated fins in adult zebrafish, likely through bioelectricity ^18^. Similarly, *kcnh2a* was pinpointed as the cause of the classical longfin mutant, *lof*, although the ectopic expression occurs in the fin fold ^15,16^. Recently, *kcnh8* was associated with naturally existing long-finned fish, the elephant ear betta fish ^48^ and *Xiphophorus* swordfish, in which *kcnh8* was highly expressed in the sword^17^. However, the mechanisms by which *kcnh2a* and *kcnh8* cause elongated fins of these fish remain to be revealed. One promising hypothesis is that the bioelectric mechanism involves multiple potassium channels, solute carriers, and connexins, as reported in zebrafish fin size mutants ^20^. Thus, it would be interesting to examine the phenotypes of *kcnh2b*, *kcnh3,* and *kcnh6a* loss-of-function mutants and their potential roles in developmental patterning in the future.

## MATERIAL AND METHOD

### Zebrafish strains, husbandry, and CRISPR

Zebrafish were raised and maintained by following AAALAC-approved standards at the Purdue animal housing facility. Zebrafish were maintained according to the Zebrafish Book ^49^. All zebrafish experiments were approved by the Purdue Animal Care and Use Committee (PACUC, #1210000750). Wildtype TAB fish (AB/Tübingen, RRID: ZIRC_ZL1) were employed for this project ^50,51^, and zebrafish embryos were staged based on the Kimmel staging guide ^52^.

### Phylogenetic and syntenic analyses

Zebrafish Kcnh channel protein sequences were first identified by searching the NCBI (RRID: SCR_006472) and Ensembl (RRID: SCR_002344) databases using human protein symbols. All zebrafish Kcnh channels with partial sequences in Ensembl were further searched for full-length sequences using BLASTp in NCBI. All the retrieved zebrafish *kcnh* genes were also validated in the ZFIN database (PRID: RRID:SCR_002560). Orthologous and paralogous genes were retrieved from Ensembl using the comparative genomics function in GRCz11. For all the orthologous protein sequences, representative protein sequences from the major metazoan taxa were selected based on their taxonomic positions and the quality of available protein sequences. When multiple isoforms were present, the longest sequence was chosen for analysis. All the protein sequences used are listed in the Supplementary **Table S1**.

Phylogenetic and syntenic analyses were performed as we described previously ^34,53–56^. Briefly, multiple protein sequences were aligned using the MUSCLE program (RRID: SCR_011812) with default settings ^57^. To identify the best evolutionary model for phylogenetic analysis, we conducted a model selection test using maximum likelihood (ML) with default parameters in MEGA12 (RRID:SCR_000667) ^58^. The models with the lowest B.I.C. (Bayesian Information Criterion) scores were considered to describe the substitution pattern the best, and JTT (Jones-Taylor-Thornton model) +G (gamma distribution) was chosen. Then, we constructed phylogenetic trees using RaxML (RRID:SCR_006086) (8.2.12) ^59^. ML phylogenetic analysis was performed using JTT + G with 1000 bootstrap replicates ^53,54^. The final phylogenetic trees were viewed and generated with FigTree (RRID:SCR_008515) V1.4.2 (http://tree.bio.ed.ac.uk/software/figtree). The syntenic analysis was first carried out in the Genomicus browser (version 100.01, RRID:SCR_011791). Individual gene positions were then verified in Ensembl, the UCSC Genome Browser (RRID: SCR_005780), and the Synteny Database ^60^.

### Zebrafish *kcnh* gene cloning

After zebrafish *kcnh* genes were identified, their coding sequences were used to design PCR primers to amplify full open reading frames (ORFs). Extra nucleotide sequences were added according to the 5’ end of the primers to facilitate In-Fusion cloning. The consensus Kozak sequences (GCCACC) were also inserted immediately before the start codon in the forward primers. All primer sequences were listed in Supplementary **Table S2**.

To extract total RNAs, about 100 embryos (1-3 days post-fertilization) were dechorionated and pooled. TRIzol reagent (Thermo Fisher, 15596026) was used for RNA extraction according to the manufacturer’s instructions. The total RNA amount and quality were checked using a Nanodrop spectrophotometer (RRID: SCR_018042). Reverse transcriptions were performed with the SuperScript® III First-Strand Synthesis System (Thermo Fisher, 18080051) following manual instructions. Phusion® High-Fidelity DNA Polymerase master mix (Thermo Scientific, #F530L) or PrimeSTAR® GXL DNA Polymerase (Takara Bio, R050A) were chosen for PCR amplification. Right-sized PCR products were examined on DNA agarose gels and purified with NucleoSpin Gel and PCR Clean-up Kit (Takara Bio, 740609.250). Purified PCR products of each gene were then cloned into the pENTR-D vector. The pENTR-D vector was prepared by PCR using pENTR-D-sp6-*kcnk4b* (RRID: Addgene #213123). Recombination cloning was performed according to the manufacturer’s instructions (Takara Bio, #638946). Correct clones were verified for correct insertional gene orientation by endonuclease digestion and nanopore sequencing. All zebrafish kchn gene clones are available from Addgene (Plasmids #255299, 255301-255313).

### Whole-mount *in situ* hybridization, cryosection, and imaging

Riboprobe DNA templates were prepared either by PCR amplification or plasmid linearization with an endonuclease (Not I) at the 5’ end of the protein-coding region in the pENTR-D vector. DNA templates were purified before *in vitro* transcription using the phenol-chloroform extraction and ethanol precipitation ^61^. Anti-sense riboprobes were synthesized by *in vitro* transcription using T7 RNA polymerase (Thermo Scientific, EP0111) and DIG RNA Labeling Mix (Millipore-Sigma, 11277073910) according to the manufacturers’ manuals. All synthesized riboprobes were purified using Sigma Spin post-reaction clean-up columns (Sigma, S5059) and stored at - 80°C in a freezer before use.

Whole-mount *in situ* hybridizations were performed according to our established method, with some modifications ^33–35^. Briefly, chorions were removed using pronase (Sigma, PRON-RO) treatment before fixation for 0.5-3 dpf fish embryos. All fish embryos were staged and then fixed with 4% PFA (paraformaldehyde) for 1-2 days at 4°C, followed by dehydration in serial methanol (25%, 50%, 75%, and 100%) with PBST (Phosphate-buffered saline solution with 0.1% tween-20). Dehydrated embryos were stored in 100% methanol at -20 °C until use. We rehydrated fish embryos using a reverse gradient of methanol in PBST (100%, 75%, 50%, and 25%) before performing *in situ* hybridization experiments. Embryos (12-24 hpf) were then bleached with 6% H_2_O_2_ in PBST and permeabilized with proteinase K (10 μg/ml in PBT). After permeabilization, embryos were washed with PBT and fixed in 4% PFA with 0.2% glutaraldehyde for at least 3 hrs at room temperature to inactivate proteinase K. Next, fish embryos were washed with PBST (2 x 10 minutes) to remove fixatives. Then, fish embryos were incubated in pre-hybridization solution [50% formamide, 5XSSC (0.75M NaCl, 75mM sodium citrate, pH 7.0.), 2% Roche blocking powder, 0.1% Triton-X, 50 mg/ml heparin, 1 mg/ml Torula yeast RNA, 1 mM EDTA, 0.1% CHAPS, DEPC-treated ddH_2_O] overnight at 65°C in hybridization oven with gentle shaking (40 rpm). Riboprobes (1µl) were added on the morning of the second day. Then, embryos were further hybridized for continuous 48-72 hours before washing unbound riboprobes with 2X SSC (Saline-sodium citrate) and 0.2X SSC (3 x 30 minutes for each solution) at 65°C. Next, fish embryos were washed with KTBT 2x for 10 minutes each (50 mM Tris-HCl, pH 7.5, 150 mM NaCl, 10 mM KCl, 1.0% Tween-20) before antigen blocking using 1% sheep serum (Sigma, S2263) and 2 mg/ml BSA in KTBT for at least 3 hours. Anti-digoxigenin antibody conjugated to alkaline phosphatase (Roche, 11093274910, RRID: AB_2734716) was added to the blocking solution at a 1:5000 dilution, and embryos were incubated overnight at 4°C with gentle shaking. After 6 times 1 hr washing with KTBT and overnight KTBT incubation at 4°C with gentle shaking, the following day, color reaction was carried out in NTMT solution (100 mM Tris-HCl, pH 9.5, 50 mM MgCl2, 100 mM NaCl, 0.5% Tween-20) with 75 mg/ml NBT, 50 mg/ml BCIP, and 10% DMF (N, N-dimethylformamide) after 2x 10-minutes NTMT wash. For 2-3 dpf fish embryos, 1mM levamisole was added to reduce the effect of endogenous alkaline phosphatase. Color development was carried out in the dark with gentle rocking. We closely monitored the color density of each reaction. Once the embryos developed suitable color densities, the reaction was stopped with NTMT washing. The finished samples were imaged immediately or stored in 4% PFA for later imaging. For histological analysis, post-hybridization embryos were equilibrated overnight in 15% sucrose, then in 30% sucrose with 20% gelatin, after which they were embedded in 20% gelatin for cryosectioning (10-25 μm) on a cryotome. Images were acquired using an Axiocam 305 color camera on a Zeiss Stereo Discovery V12 (RRID: SCR_027509) and an Axio Imager 2 compound microscope (RRID: SCR_018876). Whole-mount embryos were imaged in 3% methylcellulose.

## ACKNOWLEDGMENTS

The research was supported by the National Institute of General Medical Sciences of the National Institutes of Health (R35GM124913) to G.Z. The content is solely the responsibility of the authors and does not necessarily represent the official views of the funding agents. The authors also thank the Hayward Foundation for its generous support of the laboratory.

## AUTHOR CONTRIBUTIONS

**Kuangyi Wu** and **Dingxun Wang:** Formal analysis, Investigation, Methodology, Writing-review & editing. **Ziyu Dong** and **Alice Zhou:** Investigation, Writing-review & editing. **GuangJun Zhang:** Conceptualization, Formal analysis, Funding acquisition, Investigation, Supervision, Writing-original draft, Writing-review & editing.

## CONFLICT OF INTERESTS

The authors declare no conflict of interest.

## Notes

### Competing Interest Statement

The authors have declared no competing interest.

