## Supplementary tables for "Evolutionary analysis of vertebrate KCNH voltage-gated potassium channels and their expression in zebrafish embryos"

**Table S1. Protein sequences used for phylogenetic analysis.**

| Common species name | Scientific Name | KCNH1 | KCNH2 | KCNH3 | KCNH4 | KCNH5 | KCNH6 | KCNH7 | KCNH8 |
| --- | --- | --- | --- | --- | --- | --- | --- | --- | --- |
| Human | Homo sapiens | NP_758872.1 | NP_000229.1 | NP_036416.1 | NP_036417.1 | NP_647479.2 | NP_110406.1 | NP_150375.2 | NP_653234.2 |
| Mouse | Mus musculus | NP_034730.1 | NP_038597.2 | NP_034731.3 | NP_001074663.1 | NP_766393.2 | NP_001032801.1 | NP_573470.2 | NP_001026981.2 |
| Rat | Rattus norvegicus | XP_017454422.1 | NP_446401.1 | NP_058804.1 | NP_446082.2 | NP_598294.1 | XP_006247617.1 | NP_571987.1 | EDL83035.1 |
| Chicken | Gallus gallus | XP_040522344.1 | XP_015136636.3 |  | XP_040509293.1 | XP_040529709.1 | XP_046789286.1 | XP_046799783.1 | XP_040520526.1 |
| Turkey | Meleagris gallopavo | XP_031408030.1 | XP_010722842.2 |  |  | XP_010710334.1 | XP_010722789.1 |  | N/A |
| Xenopus | Xenopus tropicalis | KAE8600584.1 | XP_031759591.1 | XP_031753016.1 | XP_002942364.2 | XP_031746084.1 | XP_012808358.2 | XP_031749025.1 | XP_031760363.1 |
| Eastern brown snake | Pseudonaja textilis | XP_026551168.1 |  | XP_026567024.1 | XP_026566646.1 | XP_026554170.1 | XP_026573067.1 | XP_026552928.1 | XP_026564036.1 |
| Chinese softshell turtle | Pelodiscus sinensis |  |  | XP_075758072.1 | XP_075767445.1 | XP_075782488.1 | XP_075767334.1 | XP_075789622.1 | XP_006120233.2 |
| Coelacanth | Latimeria chalumnae | XP_064411209.1 | XP_014348120.1 | XP_064408372.1 | XP_064421690.1 | XP_064421237.1 | XP_014343047.1 | XP_064420119.1 | XP_064410955.1 |
| Elephant shark | Callorhinchus milii | XP_007890670.1 |  |  |  | XP_007902076.1 |  | XP_007888033.2 | XP_042188420.1 |
| Spotted gar | Lepisosteus oculatus | XP_069035919.1 | XP_006634281.2 | XP_015217397.2 | XP_015217710.2 | XP_006632393.1 | XP_015217397.2 | XP_069052888.1 | XP_069051169.1 |
| Nile tilapia | Oreochromis niloticus | kcnh1a: XP_025753448.1 | kcnh2b: XP_019218510.1 | XP_005460711.1 | kcnh4a:XP_005468906.1 | kcnh5a: XP_003449970.1 | kcnh6a: XP_005469007.1 | kcnh7a: XP_013128067.1 | XP_003448945.1 |
|  |  | kcnh1b: CAI5681500.1 |  |  | kcnh4b:XP_005471361.1 | kcnh5b: XP_003451242.1 |  | kcnh7b: XP_025758714.1 |  |
| Platyfish | Xiphophorus maculatus | kcnh1a: XP_023183743.1 | kcnh2b: XP_023182235.1 | XP_014328923.1 | kcnh4a: XP_023204390.1 | kcnh5a: XP_014324695.1 | kcnh6a: XP_014330674.1 | kcnh7a: XP_023192648.1 | XP_023200281.1 |
|  |  |  |  |  | kcnh4b: XP_023197359.1 |  |  | kcnh7b: XP_014327628.1 |  |
| Three-spined stickleback | Gasterosteus aculeatus aculeatus | kcnh1a: XP_040033853.1 | kcnh2a: ENSGACT00000053404.1 | XP_040044960.1 | kcnh4a: XP_040046504.1 | kcnh5a: XP_040017369.1 | kcnh6a: XP_040047636.1 | kcnh7a: XP_040056596.1 | XP_040045452.1 |
|  |  | kcnh1b: XP_040042776.1 | kcnh2b: XP_040022853.1 |  | kcnh4b: XP_040032699.1 |  | kcnh6b: XP_040032088.1 | kcnh7b: XP_040044756.1 |  |
| Japanese medaka HdrR | Oryzias latipes | kcnh1a: XP_020565084.1 |  |  | kcnh4a: XP_020560803.1 | kcnh5a: XP_011490256.1 | kcnh6a: XP_011476138.1 | kcnh7a: XP_011488040.1 | XP_011479155.1 |
|  |  | kcnh1b: XP_023810050.1 | kcnh2b: XP_011486965.1 |  | kcnh4b: XP_004080285.1 | kcnh5b: XP_023807694.1 |  | kcnh7b: XP_020565953.1 |  |
| Zebrafish | Danio rerio | kcnh1a: XP_073783443.1 | kcnh2a: XP_073775577.1 | XP_001919436.3 | kcnh4a: NP_001309362.1 | kcnh5a: XP_073788639.1 | kcnh6a: NP_998002.1 | kcnh7a: XP_073769333.1 | XP_073787754.1 |
|  |  | kcnh1b: XP_009294796.1 | kcnh2b: XP_073797320.1 |  | kcnh4b: XP_073774395.1 | kcnh5b: XP_002664253.1 | kcnh6b: XP_073774351.1 | kcnh7b: XP_073809787.1 |  |
| Mexican tetra | Astyanax mexicanus | kcnh1a: XP_049320079.1 | kcnh2a: XP_049323249.1 | KAG9271189.1 | kcnh4a: XP_022528303.2 | kcnh5a: KAG9282826.1 | kcnh6a: ENSAMXT00000012115.2 | kcnh7a: XP_049340940.1 | XP_049331969.1 |
|  |  | kcnh1b: XP_022525848.2 | kcnh2b: XP_007256983.3 |  | kcnh4b: ENSAMXT00000031902.1 | kcnh5b: KAG9274900.1 | kcnh6b:XP_049320999.1 | kcnh7b: XP_049335102.1 |  |
| Electric eel | Electrophorus electricus | kcnh1a: XP_035388576.1 | kcnh2a: XP_026861581.2 | XP_026851708.2 | kcnh4a: XP_035386833.1 | kcnh5a: XP_026858766.2 | kcnh6a: XP_035379849.1 | kcnh7a: XP_026866052.2 | XP_035385068.1 |
|  |  | kcnh1b: XP_026852208.2 | kcnh2b: XP_026858752.1 |  | kcnh4b: XP_026873473.2 |  |  | kcnh7b: XP_035382667.1 |  |
| Fugu | Takifugu rubripes | kcnh1a: XP_029691127.1 |  | XP_029696573.1 | kcnh4a: XP_029691913.1 | kcnh5a: XP_029705807.1 | kcnh6a: XP_011602586 | kcnh7a: XP_029703364.1 | XP_003969315.2 |
|  |  |  | kcnh2b: XP_029698764.1 |  | kcnh4b: XP_003961302.2 | kcnh5b: XP_029688754.1/ ENSTRUT00000076157.1 | kcnh6b: XP_029693733.1 | kcnh7b: XP_029695709.1 |  |
| Lancelet | Branchiostoma lanceolatum | XP_066279784.1 |  |  | CAH1249231.1 | KCNH5 | XP_066301818.1: kcnh6-like | CAH1238966.1 | XP_066263673.1 |
| Tunicate | Ciona intestinalis |  |  |  |  | XP_026694540.1 | XP_026692366.1: Kcnh6 |  | XP_009859607.1: kcnh8-like |
| Common fruit fly | Drosophila melanogaster | NP_001036275.1 | NP_477009.1: eag-like | NP_001286814.1 |  |  |  |  |  |
| Roundworm | Caenorhabditis elegans | NP_001368562.1 |  | NP_001368173.1 |  |  |  |  |  |

**Table S2. PCR primers for the zebrafish *kcnh* gene cloning.**

| Gene | Transcript ID | Primers | Sequence | PCR product size (bp) |
| --- | --- | --- | --- | --- |
| *kcnh1a* | NM_001044931.1 | Dr.kcnh1a-F | 5'-GCCCCCTTGCCACCATGGCCGGGGGACGCAGAGGACTAG-3' | 2880 |
|  |  | Dr.kcnh1a-R | 5'-CGGCGCGCCCACCCTTAGGGAACATGTCCTCCTTGTCTGTGTCTGGC-3' |  |
| *kcnh1b* | NM_001278814.1 | Dr.kcnh1b-F | 5'-GCCCCCTTGCCACCATGGCGGGGGGACGCAGAGGACTGG-3' | 2968 |
|  |  | Dr.kcnh1b-R | 5'-CGGCGCGCCCACCCTTTGAAGATATATTGTCATCCTTTTCTGAATCAGGCGAG-3' |  |
| *kcnh2a* | NM_001042722.2 | Dr.kcnh2a-F | 5'-CGCCCCCTTCACCATGCCTGTACGACGGGGACACGTTGC-3' | 3759 |
|  |  | Dr.kcnh2a-R | 5'-CGGCGCGCCCACCCTTGAGCCAGGATCAGAGGGATGTCTTTTCTGCATTG-3' |  |
| *kcnh2b* | XM_073941219.1 | Dr.kcnh2b-F | 5'-GCCCCCTTGCCACCATGCCGGTGCGAAGAGGACACGTCGC-3' | 3278 |
|  |  | Dr.kcnh2b-R | 5'-CGGCGCGCCCACCCTTGCTGCCGGGGTCTGAGCTGTGC-3' |  |
| *kcnh3* | XM_001919401.8 | Dr.kcnh3-F | 5'-GCCCCCTTGCCACCATGCCTGTGATGAGAGGTCTGCTGGC-3' | 3576 |
|  |  | Dr.kcnh3-R | 5'-CGGCGCGCCCACCCTTCAGTGGTGGTCCTTCTTCATCTATGAAGCTG-3' |  |
| *kcnh4a* | [NM_001322433.1](https://www.ncbi.nlm.nih.gov/datasets/gene/100148126/#transcripts-and-proteins) | Dr.kcnh4a-F | 5'-GCCCCCTTGCCACCATGCCGGTGATGAAAGGGCTCCTGG-3' | 3453 |
|  |  | Dr.kcnh4a-R | 5'-CGGCGCGCCCACCCTTCTGAGCGACTGTGTTTTGCTCCTCAACAG-3' |  |
| *kcnh4b* | [XM_073918295.1](https://www.ncbi.nlm.nih.gov/datasets/gene/567442/#transcripts-and-proteins) | Dr.kcnh4b-F | 5'-GCCCCCTTGCCACCATGCCAGTAATGAAGGGGCTGCTGGC-3' | 3580 |
|  |  | Dr.kcnh4b-R | 5'-CGGCGCGCCCACCCTTGTCTATTAGATCCAGGCACCAAGTGGTCTCTG-3' |  |
| *kcnh5a* | [NM_001276280.1](https://www.ncbi.nlm.nih.gov/datasets/gene/556304/#transcripts-and-proteins) | Dr.kcnh5a-F | 5'-GCCCCCTTGCCACCATGCACGGGGGAAAAAGAGGACTGGTG-3' | 3279 |
|  |  | Dr.kcnh5a-R | 5'-CGGCGCGCCCACCCTTAAGCGGCCCATCATCGCCCTCCATC-3' |  |
| *kcnh5b* | [XM_002664207.7](https://www.ncbi.nlm.nih.gov/datasets/gene/100331767/#transcripts-and-proteins) | Dr.kcnh5b-F | 5'-GCCCCCTTGCCACCATGCCCGGGGGAAAGAGAGGGCTGG-3' | 3015 |
|  |  | Dr.kcnh5b-R | 5'-CGGCGCGCCCACCCTTTGGCAAGCCCTCATCTTTTTCAGAGTCCGG-3' |  |
| *kcnh6a* | [NM_212837.1](https://www.ncbi.nlm.nih.gov/datasets/gene/405763/#transcripts-and-proteins) | Dr.kcnh6a-F | 5'-GCCCCCTTGCCACCATGCCCGTGCGCCGCGGACATGTC-3' | 3555 |
|  |  | Dr.kcnh6a-R | 5'-CGGCGCGCCCACCCTTGCTTCCGGGTAAGACTGGATCGGACACG-3' |  |
| *kcnh6b* | [XM_021472507.3](https://www.ncbi.nlm.nih.gov/datasets/gene/570070/#transcripts-and-proteins) | Dr.kcnh6b-F | 5'-GCCCCCTTGCCACCATGCAGCGCTTCAACGGCTTGAAGAGC-3' | 2679 |
|  |  | Dr.kcnh6b-R | 5'-CGGCGCGCCCACCCTTGCTGACCGGTAATCCAGGATCAGACATGTG-3' |  |
| *kcnh7a* | [XM_073953689.1](https://www.ncbi.nlm.nih.gov/datasets/gene/560289/#transcripts-and-proteins) | Dr.kcnh7a-F | 5'-GCCCCCTTGCCACCATGCCTGTGCGGAGAGGTCATGTC-3' | 3405 |
|  |  | Dr.kcnh7a-R | 5'-CGGCGCGCCCACCCTTGTGCTGCTGGTCGGGTGGTCC-3' |  |
| *kcnh7b* | XM_073913232.1 | Dr.kcnh7b-F | 5'-GCCCCCTTGCCACCATGCCGGTGCGACGGGGTCACGTC-3' | 3531 |
|  |  | Dr.kcnh7b-R | 5'-CGGCGCGCCCACCCTTCTGCCCTGGAAGGCCGGGATCAGAG-3' |  |
| *kcnh8* | XM_068215273.2 | Dr.kcnh8-F | 5'-GCCCCCTTGCCACCATGCCTGTTATGAAGGGATTGCTCGCTCCAC-3' | 2988 |
|  |  | Dr.kcnh8-R | 5'-CGGCGCGCCCACCCTTCAAACCTAGGGACCTGAGATCCTCCCCTTC-3' |  |
| Note: In-fusion cloning adapter sequences are derlined | | | | |
